# A Fast Likelihood Method to Reconstruct and Visualize Ancestral Scenarios

**DOI:** 10.1101/379529

**Authors:** Sohta A. Ishikawa, Anna Zhukova, Wataru Iwasaki, Olivier Gascuel

**Affiliations:** Unité Bioinformatique Evolutive, C3BI USR 3756, Institut Pasteur & CNRS, Paris, France; Department of Biological Sciences, the University of Tokyo, Tokyo, Japan; Méthodes et Algorithmes pour la Bioinformatique, IBC - LIRMM UMR 5506, CNRS & Université de Montpellier, Montpellier, France

**Author notes:** Joint first authors (Sohta A. Ishikawa, Anna Zhukova).

**Keywords:** phylogenetics, ancestral character reconstruction, maximum likelihood, marginal and joint posterior probabilities, maximum a posteriori, Brier scoring rule, simulations, HIV, phylogeography, drug resistance mutations

## Abstract

The reconstruction of ancestral scenarios is widely used to study the evolution of characters along a phylogenetic tree. In the likelihood framework one commonly uses the marginal posterior probabilities of the character states, and the joint reconstruction of the most likely scenario. Both approaches are somewhat unsatisfactory. Marginal reconstructions provide users with state probabilities, but these are difficult to interpret and visualize, while joint reconstructions select a unique state for every tree node and thus do not reflect the uncertainty of inferences.

We propose a simple and fast approach, which is in between these two extremes. We use decision-theory concepts and the Brier criterion to associate each node in the tree to a set of likely states. A unique state is predicted in the tree regions with low uncertainty, while several states are predicted in the uncertain regions, typically around the tree root. To visualize the results, we cluster the neighboring nodes associated to the same states and use graph visualization tools. The method is implemented in the PastML program and web server.

The results on simulated data consistently show the accuracy and robustness of the approach. The method is applied to large tree comprising 3,619 sequences from HIV-1M subtype C sampled worldwide, which is processed in a few minutes. Results are very convincing: we retrieve and visualize the main transmission routes of HIV-1C; we demonstrate that drug resistance mutations mostly emerge independently under treatment pressure, but some resistance clusters are found, corresponding to transmissions among untreated patients.

## Introduction

A central issue in biology is to recover and understand the evolutionary history of biological entities. These may be of different nature and scale, ranging from DNA and proteins to communities, going through biological systems, organs, strains, individuals, species and populations. The characteristics and evolution of these objects are measured using a variety of “characters”, including molecular properties (e.g, Werner et al. 2014, Bickelmann et al. 2015, Busch et al. 2016), gene-contents of genomes (e.g, Iwasaki and Takagi 2007), morphological and phenotypic characteristics (e.g, Endress and Doyle 2009, Marazzi et al. 2012, Beaulieu et al. 2013, Sauquet et al. 2017), ecological traits (e.g, Maor et al. 2017), and geographic locations (e.g. Wallace et al. 2007, Arbogast 2001, Lemey et al. 2009, Edwards et al. 2011, Lemey et al. 2014, Magee et al. 2017, Dudas et al. 2017). Ancestral character reconstruction (ACR) is a major approach to tackle these questions, allowing us to trace the origin and evolution of the character of interest. ACR relies first on the inference of phylogenetic relationships among the studied objects, that is, a phylogenetic tree. Tips of the tree represent extant objects, progressively connected by branches to their ancestors represented by the internal nodes of the tree. The common ancestor of all tips corresponds to the tree root. ACR determines how the character has changed on the tree from the root to the tips over evolutionary time, by assigning the most likely ancestral character states to each internal node. This global reconstruction over the whole tree describes the evolutionary history of the character of interest and is commonly called an “ancestral scenario”. Several approaches have been proposed for ACR so far, including parsimony (Swofford and Maddison 1987, Maddison and Maddison 2000), maximum-likelihood (ML; Felsenstein 1981, Pagel 1999, Pupko et al. 2000, Ree and Smith 2008) and Bayesian methods (Huelsenbeck and Bollback 2001, Pagel et al. 2004).

Parsimony-based ACR provides quick and simple methods to infer ancestral scenarios. However, due to the over-simplification of evolutionary processes (e.g. not accounting for branch lengths and evolutionary times), parsimony has limited accuracy (Zhang and Nei 1997, Collins et al. 1994). ML and Bayesian approaches are based on probabilistic models of character evolution. ML methods were shown to perform better than parsimony, using both theoretical arguments and simulation studies under a variety of conditions (Zhang and Nei 1997, Gascuel and Steel 2014). Simulation results showed that even the simplest models (e.g. JC, Jukes and Cantor 1969) yield more accurate reconstructions than parsimony (Gascuel and Steel 2014), thanks to the consideration of evolutionary times and branch lengths, and are robust to moderate model violations and phylogenetic uncertainty (Hanson-Smith et al. 2010).

The size of the trees subjected to ACR has rapidly increased thanks to new generation sequencing technologies. Evolutionary and epidemiological analysis of pathogens like human immunodeficiency virus (HIV), Influenza and Ebola is one of the hotspots of this problem, with data sets commonly comprising thousands strains (Ratmann et al. 2016, Holmes et al. 2016, Durães-Carvalho and Salemi 2018). With such rapidly evolving pathogens, the links between evolutionary and epidemiological processes raise essential public health questions with important practical issues, notably the routes and patterns of pathogen spread (Gräf et al. 2015) and the emergence of drug resistances (Mourad et al. 2015). ACR has been widely applied to tackle these questions aiming to map ancestral states of pathogen characters (e.g. sampling location, risk group of the host, presence of drug resistance) on the tree inferred from genetic sequences.

Bayesian methods (Huelsenbeck and Bollback 2001, Pagel et al. 2004, Drummond et al. 2012) are commonly used in this context, notably in phylogeography studies (Lewis et al. 2015, Magree et al. 2017). The main approach is to infer using a Markov chain Monte Carlo (MCMC) procedure the joint posterior distribution of ancestral character states, phylogenetic tree and model parameters. This involves complex probabilistic models describing the evolution of the sequences, the molecular clock (possibly relaxed and correlated), the demography, and last but not least the evolution of the studied character. The character evolution model can be very simple, typically symmetrical with a few states, but the current trend is to rely on increasingly complex models, non-symmetrical, with latent variables, dozens of character states, and evolution over time (Stadler and Bonhoeffer 2013, Leventhal et al. 2013, Kühnert et al. 2014, Lambert et al. 2014, Kühnert et al. 2016). The Bayesian approach is very popular because of this wealth of options and flexibility, via famous software programs like BEAST (Drummond et al. 2012). However, MCMC-based methods have high computational cost, and the joint inference of all these tree, parameter and character distributions cannot be achieved for large data sets. Even the stepwise approach where we first infer the tree distribution, and then the distribution of the studied character along the most likely trees is hardly applicable to big datasets, requiring high performance computing units (typically GPUs) and sophisticated parallel implementations (Ayres et al. 2011). In contrast, the ML approach is less computationally demanding as it gives point estimates for the parameters of interest, instead of distributions. For example, TreeTime (Sagulenko et al. 2018) is able to deal with large trees with thousands of tips and perform fast ML-based ACR in a few minutes or even a few seconds.

However, there are still potential limitations in applying standard ML-based ACR to large datasets and trees. These limitations are related to the inference of the character states, the uncertainty that is inherent to such inference and that of the phylogeny, and the visualization and interpretation of the (large) resulting ancestral scenario. Two main approaches are used in ML-based ACR:

- Either we compute the marginal posterior probabilities of every state for each of the tree nodes (Felsenstein 1981, Yang 2007). Then, we usually select the state with the highest posterior. This maximum a posteriori (MAP) selection is independent from one node to another, which could induce globally inconsistent scenarios (typically: two very close nodes with incompatible predictions). This possible shortcoming formed the basis of criticisms against the marginal approach (but see Gascuel and Steel (2014) and our simulation results below).
- Or we compute with dynamic programming the joint ancestral scenario with the maximal posterior probability (Pupko et al. 2000). This approach has some global consistency guaranty, but does not reflect the fact that with real data and large trees, billions of scenarios may have similar posterior probabilities.

From a theoretical standpoint, both methods are based on MAP and thus have some optimality guaranty, at least in the absence of model violation. Simulations performed by Gascuel and Steel (2014) showed that the predictions and accuracy of both are extremely close. This advocates for the use of the marginal approach, which not only indicates the most likely state for each node, but also returns the posteriors of all states. However, interpreting and using these probabilistic outputs is difficult, for example when two states have similar posteriors. Another difficulty is to visualize and summarize the resulting, global scenario, which commonly involves thousands of probability distributions attached to each of the tree nodes.

Here, we propose a simple and fast approach to overcome these limitations. We use decision theoretic concepts and tools to infer for each of the tree nodes a limited set of likely states, which best approximate the marginal posterior probabilities. In the easy regions of the tree (typically close to the tips, Gascuel and Steel 2014) this approach predicts a unique (MAP) state, while in the difficult parts (typically close to the root) it may predict several likely states reflecting the uncertainty of the inferences. To summarize and visualize the results we cluster the neighboring nodes with identical predictions and re-use some of the ideas we developed in parsimony-based PhyloType software (Chevenet et al. 2013). Thus we obtain a compact, tree-shaped and easily interpretable graphical representation of the most likely ancestral scenarios, which is robust to phylogenetic uncertainties and sampling rate variations. In the following, we first describe the different components of the method, then the results with simulated data along with comparisons with other ACR methods, and lastly the analysis of a large HIV data set. All methods developed and studied in this article are implemented in PastML software, which is freely available in several versions and interfaces (open source code, docker container, web server, see https://pastml.pasteur.fr/).

## New Approaches

### Preamble

The method can be decomposed into three main steps: (1) ML-based rescaling of the tree and estimation of the model parameters; (2) ancestral reconstruction of the most likely character states; (3) compression and visualization of the inferred ancestral scenario. These three steps are described in turn in the following. In this section, we describe the input data, notation, model and global framework and goals.

The input of the method is a rooted tree denoted as *T*, where every tip is associated to a character state. The number of tree tips is denoted as *n* and the tree root as *R. T* may be not fully resolved, the method applies to both binary and non-binary trees. In most cases *T* is obtained from a multiple alignment of sequences (DNA or proteins) using some standard phylogenetic software. Then, the branch lengths are expressed in number of substitutions per site. As we shall see, the input tree is rescaled to fit the evolution of the studied character, and thus all branch-length measures are acceptable. Most interesting results will be obtained with time scaled trees, where branch lengths are expressed in years. Then, the rescaling factor estimated from the input data represents the average number of character changes per year. When the goal is to reconstruct the ancestral sequences corresponding to the input multiple alignment, rescaling is not needed and the original branch lengths in the tree are well suited.

The studied character may be of various nature, as discussed in the Introduction. Here, we consider discrete characters with values taken from a finite, non-ordered set of states; for example: {A, T, G, C} for DNA, {Africa, America, Asia, Australia, Europe} in phylogeography, or {Sensitive, Resistant} when studying drug resistances. *S* denotes the set of possible states, with size 5. A tree tip (or leaf) is denoted as l, and *c (l)* ∈ *S* is the character state associated to *l*. The method is able to accommodate tips with unknown character values, denoted as *c* (*l*) = *X*.

Continuous-time Markov models are commonly used to represent the evolution of characters, notably with sequences where all multiple-alignment sites are usually assumed to evolve according to the same model (with different rates when using rates across sites models, e.g. gamma distributed, Yang 1994). In this setting, especially with DNA where we have 4 states only, we are able to accurately estimate the parameters of relatively complex models, for example GTR (Tavaré 1986) having 10 parameters and 8 degrees of freedom with DNA. Here, we have a unique observation describing the evolution of the studied character through the tips values. Then, accurately estimating the parameters of complex models is a difficult task, which may just be impossible, especially when *S* is large. We thus use simple JC-like and F81-like models (Jukes and Cantor 1969, Felsenstein 1981). With JC-like models all rates of changes from state *i* to state *j* (*i ≠ j*) are equal, while with F81-like models, the rate of changes from *i* to *j* (*i ≠ j*) is proportional to the equilibrium frequency of *j*, denoted as *π j*. JC-like models are special cases of F81-like ones, with all equilibrium frequencies equal to 1/ 5. Several studies advocate the use of F81-like models. As said above, we showed using simulations that even the simpler JC version performs nearly as well as the true model, with DNA data generated using an HKY model (Hasegawa et al. 1985). Moreover, Dudas et al. (2017) showed that the main factor of state changes in Ebola phylogeography (i.e. virus dispersal) corresponds to a number of movements from location *i* to location *j* proportional to the product of their population sizes *π_i_π_j_*, which precisely corresponds to an F81-like model. Another advantage of F81 is that the probability of changes along a branch of length *t* is simply expressed as:

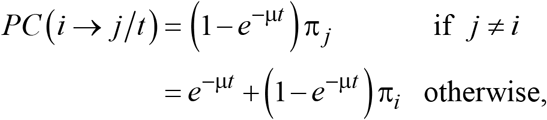

where μ is the normalization factor:

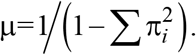

F81-like models (also called equal input models) have 5 parameters (5-1 degrees of freedom) corresponding to the equilibrium frequencies of the 5 states. In our software, these frequencies can be user supplied, roughly estimated from the state frequencies observed at the tree tips (not recommended), or estimated by maximum likelihood as we shall see in the next section.

### Tree rescaling and parameter estimation

Beyond the state equilibrium frequencies, the whole model involves two additional parameters:

- The global rate, denoted as p. With a unique observation, as is the case here, estimating all branch lengths in the tree is just impossible. We therefore assume that the number of character changes along the tree is proportional to the branch lengths of the input tree. It follows that every branch length *t* is turned into *ρt*, which is interpreted as the expected number of character changes along the given branch. Moreover, we assume that *ρ* is constant across the tree over evolutionary time, which is a similar assumption as the one-rate model (Mooers and Schluter 1999). With dated input trees, the original branch lengths are measured in years and *ρ* in number of state changes per year. The estimated value of p is then highly informative about the global evolutionary rate of the studied character along the tree.
- A smoothing parameter, denoted as ε. Both dated and molecular trees may have branches of length zero. For example, when two input sequences are identical (quite common with virus strains), we expect that any reasonable phylogenetic method infers a cherry with null branches connecting the two sequences (a cherry is a rooted subtree of two taxa). The same configuration may happen in dated trees due to temporal constraints (To et al. 2015). However, two identical sequences may have been observed in different countries, thus giving rise to two different character states linked by a path of length zero. The likelihood of any scenario containing such a configuration is null and no ML-based ancestral reconstruction is possible. Thus, we use a smoothing approach to lengthen null and short external branches, without modifying their average length, denoted as *τ*. Let *t* be the length of a given external branch of the input tree, the smoothed length is equal to *τ*(*t + ε*))(*τ + ε*). Zero length internal branches are not smoothed but turned into polytomies, as only null external branches may be problematic. The combination of the global rate with the smoothing procedure provides the “rescaled” tree with external branches of length ρτ(*t* + ε)/(τ + ε) and internal ones of length ρ*t*, where *t* is the original length in the input tree.

To estimate these parameters (ρ, ε, and the equilibrium frequencies with F81-like models) we compute the scenario likelihood using the standard pruning algorithm (Felsenstein 1981) and optimize this likelihood using the iterative Broyden–Fletcher–Goldfarb–Shanno (BFGS) algorithm (Fletcher 2013). Constraints are added to account for the biological/mathematical meaning of the parameters:

- Let β be the average branch length in the input tree, excluding branches with length zero. We impose: β^−1^ x 1/1000 ≤ ρ ≤ β^−1^ x10, meaning that in the rescaled tree the number of changes along a branch with average length β is in-between 0.001 and 10 (note that with 10 changes per branch reconstructing ancestral scenarios is just impossible).
- Let τ be the average length of external branches in the input tree, excluding branches with length zero. We impose τ/100 ≤ ε ≤ τ/10, meaning that null branches will have length in-between ~ τ/100 and ~ τ/10, before rescaling, while longer branches will not change substantially.
- Last but not least, we (obviously) impose Σ π_*i*_ = 1.

To ensure that these constraints are satisfied, we use variable transformations: softmax for the equilibrium frequencies, and sigmoid-based for the scaling factor ρ and smoothing parameter ε. The corresponding variables are then optimized in the full multidimensional space using (unconstrained) BFGS.

### Discrete approximation of the state marginal posterior probabilities

Our method is based on a discrete approximation of the marginal posterior probabilities of the character states, attached to the internal nodes of the tree. The computation of these probabilities is standard and used under different forms in most if not all ML-based phylogenetic programs. However, the complete description of the procedure is rarely available, and not presented in any text book to the best of our knowledge. It is described in the Material and Methods, for the sake of completeness. To summarize: we first use the pruning algorithm (Felsenstein 1981), which performs a bottom up, post-order tree traversal, and accounts for the information of the descendants of every tree node; then, we perform a top-down, pre-order tree traversal, which adds to the previous calculations the information coming from the rest of the tree. We thus obtain for every tree node *N* and state i, the marginal posterior probability of *i* for *N, Marginal*(*N,i*), which accounts for the state value of all tree tips. This procedure has a time complexity in *O* (*ns*^2^), where *n* is the number of tips and s the number of states. It is thus linear in *n* and able to process trees with dozens of thousands of tips in a few seconds. It is equivalent (but faster) to the procedure consisting in iteratively re-rooting the tree with every internal node and applying the pruning algorithm. The reconstruction accuracy is clearly higher than that obtained with the pruning algorithm (without re-rooting) and the descendant information only (Gascuel and Steel 2014).

Let *N* be any given internal node of *T*. Based on the marginal posterior probabilities *Marginal* (*N, i*), we have to decide which states are predicted for *N* and which ones are discarded because their posteriors are too low. We could use some thresholding approach, but the choice of the threshold values and decision procedure would be very subjective, without any formal guaranty on the accuracy of the predictions. We therefore used concepts and tools from decision theory and supervised classification (Brier 1950, Gneiting and Raftery 2007).

Assume that the true evolutionary model (tree, branch lengths, model of character changes) is fully known; then, a standard result, known as the Bayes decision rule, is that the most accurate prediction for *N* is obtained by selecting the state with highest posterior (MAP, maximum a posteriori). In this framework, we predict a unique state, and the accuracy is simply measured by the probability of correctly predicting the true ancestral state. However, in our framework a unique prediction per node is often unsatisfactory, especially when several states corresponding to different scenarios have similar posteriors. Then, a refined approach involves using probabilistic predictions, where states are assigned to probabilities instead of binary, mutually exclusive decisions as with the Bayes rule. Again, when the true evolutionary model is fully known, the marginal posterior probabilities can be shown to be optimal. Various scoring criteria (or scoring rules) have been proposed to measure the accuracy of probabilistic predictors. The most used is the logarithmic scoring criterion, which has strong foundations in information theory. This is the negative of “surprisal”, which is commonly used in Bayesian inference. However, this scoring criterion is not appropriate in our context, where we have state predictions with null probabilities (see below). We thus use the Brier quadratic scoring criterion. Let *PPr* (*N, i*) be the predicted probability of state *i* for node *N* and *Truth* (*N, i*) be the “truth” of *i* for *N*, which is equal to 1 when the ancestral state of *N* is i, and 0 otherwise. The Brier score can be expressed as:

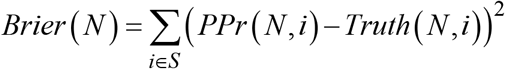

In this form the Brier score is simply the squared Euclidean distance between *PPr* (*N*) and *Truth* (*N*) (the lower the better). For instance, assuming that we assign probability 1 to the true state, then *Brier* (*N*) = 0. On the opposite, if we assign probability 1 to an incorrect state, then *Brier* (*N*) = 2, which is the worst possible value of the score. Assume now that we have no information on the ancestral state of *N*. Then, we have two natural solutions: (i) assign probability *1/s* to every state, then *Brier*(*N*)=(1 -1/*s*) +(*s*-1)(1/*s*) = 1 -1/*s*; (ii) randomly, uniformly predict one of the states, then the expected value of *Brier* (*N*)is equal to 0 ×1/s + 2 ×(*s* -1)/*s* = 2 – 2/*s*. In other words, random predictions are worse than the recognition of our ignorance.

As already said, when the model is fully known, predicting the marginal posterior probabilities of the states is optimal, regarding the Brier criterion (and other proper scoring rules, as the logarithmic one). We thus use a discrete approximation of the posteriors, which is consistently selected using the Euclidean distance. The goal is to add as little as possible error to the (unknown) optimal value of *Brier* (*N*). Assuming that we decide to retain *k* states (among 5) in the predictions, then each of these has probability 1/*k*, while the discarded states have probability 0. These probabilities are used in the selection of state subsets, but are implicit. The method returns a set of likely states without any associated probabilities. To define the state subsets to be explored, we rank the states based on their posteriors: *i*_1_ (=MAP) is best and *i*_5_ has the lowest posterior. Then, we select the best subset *SS_k_* ={*i*_1_, *i*_2_. *i_k_*} (*k*=1 *to s*) by minimizing the Euclidean distance between *Marginal* (*N*) and the probability vector defined by Pr (*i*_1→*k*_)=1/*k* and Pr(*i*_*k*+1→*s*_)=0.

This method is both simple and fast, with time complexity in *O* (*ns*^2^) again. Its accuracy strongly depends on the accuracy of the marginal posteriors, and thus on the severity of model violations, which are inevitable with real data. A possible shortcoming could be that these computations are performed independently for each of the nodes. However, we observed with simulations (Gascuel and Steel 2014) that joint reconstructions have no advantage over marginal ones, likely due to the fact that the conditional likelihoods of neighboring nodes are strongly dependent (see formulae in Methods section). Moreover, we tested a number of more sophisticated (and time consuming) methods to guaranty that node predictions are globally consistent, but did not observe any superiority over this simple approach (data not shown).

### Tree compression and visualization

On large phylogenies with hundreds or thousands of tips, once the ancestral states are reconstructed on each node, it might be difficult for a human eye to visualize and interpret the result. To overcome this issue we provide a compressed representation of the ancestral scenarios, which highlights the main facts and hides minor details. This representation is calculated in two steps: (i) “vertical merge” that clusters together the parts of the tree where no state change happens, and (ii) “horizontal merge” that clusters independent events of the same kind. Algorithmically, the two merges are performed in the following way:

- Vertical merge (vertical arrow in Figure 1): while there exists a parent-child couple such that the parent’s set of predicted states is the same as the child’s one, merge them. Moreover, we compute the size of so-obtained clusters (i.e. nodes in the compressed tree or “phylotypes” as named by Chevenet et al. (2013) in a parsimony framework) by the number of tips of the initial tree contained in it, as the tips correspond to the input data units used for tree and ancestral scenario reconstructions. Accordingly, in the initial tree each tip has a size of 1, each internal node has a size of 0, and when merging two nodes we sum their sizes.
- Horizontal merge (horizontal arrow in Figure 1): starting at the root and going top-down towards the tips, at each node we compare its child subtrees. If two or more identical subtrees are found, we keep just one representative and assign their number to the size of the branch that connects the kept subtree to the current node. Hence a branch size corresponds to the number of times its subtree is found in the initial tree. Before the horizontal merge all branches have size 1. Two trees are considered *identical* if they have the same topology and their corresponding nodes have the same state(s) and sizes. For example, two trees are identical if they are both cherries (a parent node with two tip children) with a parent in state A of size 3, a child in state B of size 2, and the other child in state C of size 1. However, those trees will not be identical to any non-cherry tree, or to a cherry with a parent not in state A, or to a cherry with the B-child of size 6, etc.

**Figure 1:**
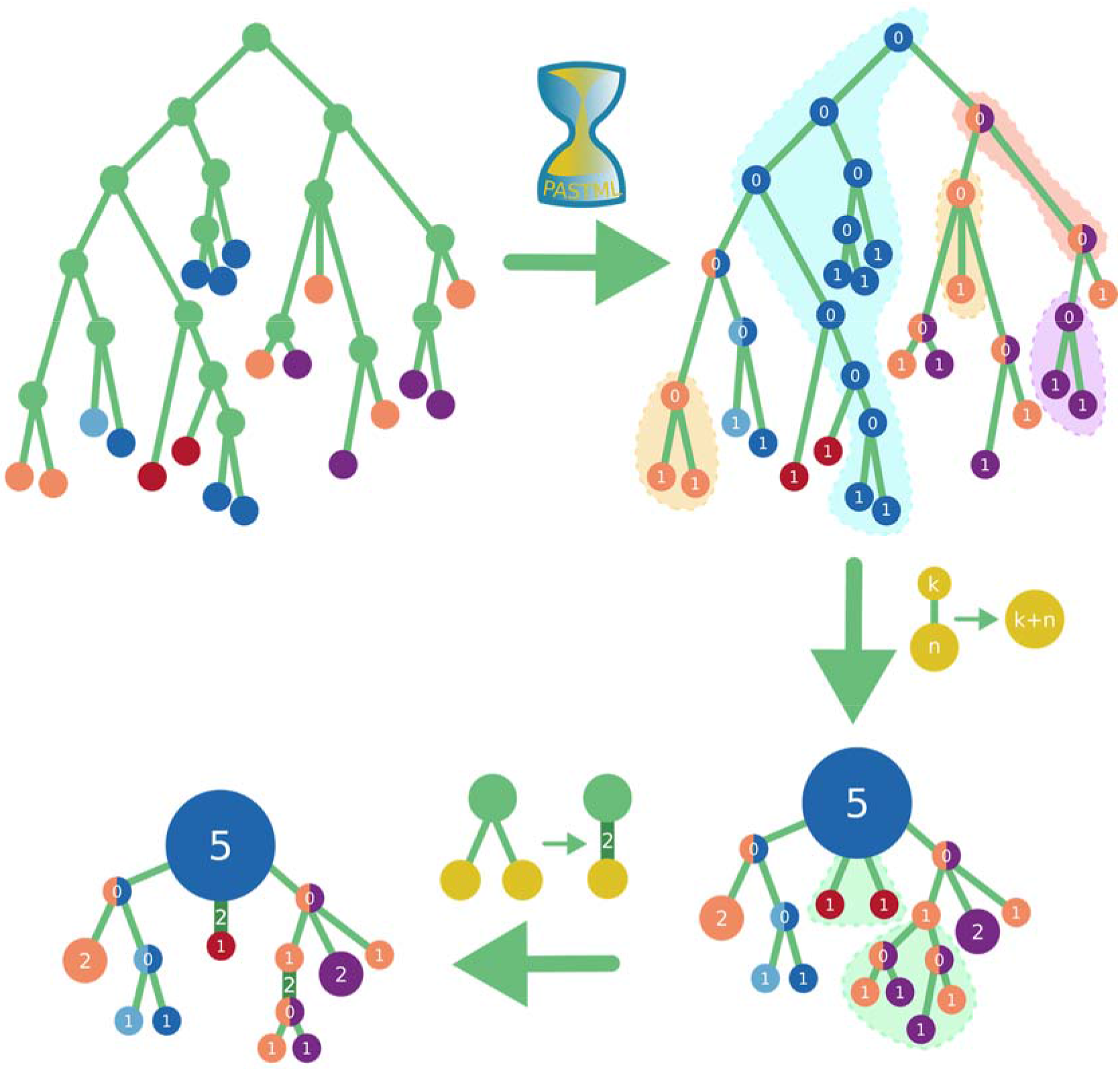
Ancestral state reconstruction and visualization steps. Starting from the initial tree with annotated tips (top left, different annotations correspond to different colors), we reconstruct the ancestral node states (top right; colored sectors are used for ambiguous nodes, e.g. green and blue), and then perform a two-step compression: the vertical compression (bottom right) clusters together the regions of the tree where no state change happens and puts the number of tips collapsed into each node as its size (e.g. the blue root cluster of size 5 in the bottom right tree corresponds to the part of the top-right tree highlighted blue and containing 5 tips), while the horizontal compression (bottom left) merges identical subtree configurations, keeping their number as branch sizes (e.g. the two red tip children of the bottom right root got merged into a red tip attached with a branch of size 2 in the bottom left tree).

These two routines are illustrated in Figure 1. In the case of a transmission tree with states representing countries, the vertical merge will cluster together the transmissions happening within the same country and having the same source within that country; for instance, see the clouds colored in blue, orange and purple in Figure 1. This operation is closely related to phylotyping (Chevenet et al. 2013), where a “phylotype” is a set of tips (strains) having the same state, as well as their most recent common ancestor (MRCA) and all nodes along the path from the MRCA to the phylotype tips (hence the vertical merge procedure). The main difference is that here we may have internal clusters with no tips, corresponding to uncertain nodes with 2 or more predicted states. Then, the horizontal merge detects independent transmissions from a country A to a country B; for instance in Figure 1, the two red nodes (= B) that branch independently from a big blue circle (= A).

For large trees with many state changes even after the compression the visualization might contain too many details. To address this issue, a program option makes it possible to relax the definition of identical trees for the horizontal merge: instead of requiring identical sizes of the corresponding nodes, we allow for nodes of sizes of the same order (log10); for instance, now a node in state A of size 3 can correspond to a node in state A of any size between 1 and 9, and a node in state B of size 25 can correspond to a node in state B of any size between 10 and 99.

If even after a relaxed horizontal merge the compressed representation contains too many details, an additional option removes minor details using the following procedure. For each leaf node we calculate its importance by multiplying its size by the size of its branch; for instance, a leaf of size 2 with a branch of size 3 gets an importance of 6. We then set the cutoff threshold to the 15-th largest node importance (a parameter that can be adjusted), and iteratively remove all leafs with smaller importance. Finally, we rerun horizontal merge as some of the previously different topologies might become identical after trimming.

To simplify the ancestral state analysis of phylogenetic trees with multiple character data, we developed a pipeline that combines the results for different characters, for example geographical location and resistance to drugs. The user provides as input the tree and a table containing the state values for the tree tips. This table is horizontally indexed by the tip identifiers, with columns corresponding to the characters. The user can choose the characters to be analyzed. We then apply ancestral reconstruction separately for each character to obtain their ancestral states, and visualize each character on the tree nodes as sectors (see application to HIV below). If we could not choose a unique state for a character, we keep the corresponding sector uncolored (i.e. white). Once the tree is colored and each node is assigned its combined states (pie of colors), we compress the tree as described in the previous section.

When sampling dates are available for the tree tips, we also provide an option to visualize a timeline: For each year between the year of the first sampled tip and the year of the last sampled one, we prune the tree to remove the tips sampled after this year, and add a slider to the visualization, allowing to navigate in time.

### Software and utilities

PastML takes as input a rooted tree and a tip state annotation table. It produces a table with predicted ancestral states, and an interactively modifiable visualization (an html file that can be viewed in a browser). PastML is available in several versions and interfaces:

- The ancestral character reconstruction algorithms discussed in this article are implemented in C, with the GNU scientific library version of the BFGS algorithm. This includes: our new Marginal Posterior Probabilities Approximation (MPPA) algorithm; the standard marginal posterior probability approach (both MAP and full probabilistic predictions); the joint posterior probability estimation algorithm of Pupko et al. (2000); and the three usual variants of parsimony-based ACR: ACCTRAN, DELTRAN, and DOWNPASS (Swofford and Maddison 1987, Maddison and Maddison 2000). Our code is open source and available from https://pastml.pasteur.fr.
- The visualization and compression procedures are implemented in Python 3.6 and JavaScript, using the Cytoscape.js library (Franz et al. 2016) for tree visualization. We implemented a Python 3 wrapper of the core C PastML library to allow for a seamless use of the whole functionality within Python 3 or from a command line. The package is available on pip3 as CytoPast. The source code and examples are available from https://pastml.pasteur.fr. We also provide a docker container that includes all the functionality and does not require installing python/C libraries: evolbioinfo/pastml.
- Last but not least, a user friendly web application is available to perform ACR, visualization, and online edition of ancestral scenarios: https://pastml.pasteur.fr.

## Results: Method Comparison Using Simulated Data

### Simulation protocol

In this study, we basically followed the simulation procedure used in (Gascuel and Steel 2014). We generated pure-birth trees with *n* = 1,000 tips. To obtain a broad range of ACR difficulties, we used different values of the speciation/substitution rate ratio (ω), which was equal to 0.1, 0.2, 0.3, 0.4, 0.5, 0.6, 0.7, 0.8, 0.9, 1.0, 2.0, 3.0, 4.0, 5.0, 6.0, 7.0, 8.0, 9.0, and 10.0. In other words, the average number of substitutions per branch was respectively equal to 5.0, 2.5, 1.67, 1.25, 1.0, 0.83, 0.71, 0,625, 0.56, 0.5, 0.25, 0.167, 0.125, 0.1, 0.083, 0.071, 0. 0625, 0.056, and 0.05 (Steel and Mooers 2010). With a high number of substitutions per branch (e.g. 5.0) ACR is very difficult, especially for the tree root, while with a low number of substitutions (e.g. 0.05) ACR becomes easy as all tips and nodes tend to have the same state value. Fifty trees were generated for each value of ω. We simulated both DNA and protein data sets along these trees, using Seq-Gen v1.3.2 (Rambaut and Grassly 1997). For DNA, nucleotide sequences of 50 sites were generated using the HKY model (Hasegawa et al. 1985) with equilibrium frequencies of A, C, G and T being equal to 0.2, 0.1, 0.3, and 0.4, respectively, and a transition/transversion ratio of 8.0. These relatively extreme values were chosen to challenge ACR when using the F81 and JC models, as implemented in PastML. Likewise, protein sequences of 50 sites were generated using the JTT model (Jones et al. 1992) with its default amino-acid equilibrium frequencies. We thus obtained 19 (ω values) x 50 (1000-tip trees) x 50 (sequence length) x 2 (DNA/protein) datasets to assess the accuracy of ACR methods. During the Monte-Carlo simulation procedure with Seq-Gen, we memorized the ancestral states of the nucleotide/amino-acid character seen at each internal nodes, including the root. Thus, the ‘true’ ancestral scenario was known. All these data are available from https://pastml.pasteur.fr/.

### Methods being compared

Simulated trees and tip state values were then subjected to ACR with five different methods:

- Parsimony: We computed the most parsimonious states for all nodes including the root. This computation was performed by two tree traversals (Maddison and Maddison 2000), analogous to the calculation of marginal posteriors (see Methods). The first post-order, bottom-up (Fitch-Hartigan or UPPASS) algorithm assigns to each parent node the most parsimonious state(s) based on the states of its child nodes. The second pre-order, top down (DOWNPASS) algorithm combines for each node the state information from its children and parent (i.e. the information derived from all tree tips). The DOWNPASS algorithm provides all most parsimonious states for all nodes, as opposed to ACCTRAN and DELTRAN heuristics which have been designed to solve ancestral ambiguities. This method returns a set of possible states for each node, and thus shares some common points with MPPA.
- Joint: we used the dynamic programming algorithm described in (Pupko et al. 2000) to infer the most likely ancestral scenario over all the tree and possible state values. For each of the nodes we thus obtained a joint estimation of the most likely state. This method returns a unique state for each node.
- Marginal: we computed the marginal likelihoods and posterior probabilities of all states for all internal nodes including the root (see Methods). This method returns full probabilistic predictions (each state is assigned a probability) for all nodes. Marginal is the best possible probabilistic predictor when the model is fully known.
- Maximum-a-Posteriori (MAP): Using previous computations, we assigned to each node the state having the highest marginal posterior. MAP thus returns a unique state for each of the nodes, just as Joint.
- Marginal Posterior Probabilities Approximation (MPPA): we used above described method to approximate the state posteriors and return for every node a subset of likely states minimizing the prediction error measured by the Brier score.

With the ML-based methods (i.e. Joint, Marginal, MAP and MPPA), the simulated datasets were analyzed under the six following conditions, corresponding to various model violations intended to measure the robustness of the ACR methods being compared:

- True model and branch lengths: the true substitution model (i.e. the model used in simulations, HKY for DNA and JTT for proteins) was used for ACR. Moreover, the model parameters were set to the same values as used in simulations, and we did not rescale the tree, keeping the branch lengths equal to those used to generate the data. Smoothing was useless as no branch had length zero, due to the tree generation procedure. In this setting we had no model violation and the accuracy was the best possible. The goal was to check that the results were only slightly degraded in the other conditions, which include various levels of model violation.
- True model and flattened branch lengths: just as in the previous condition we used the true substitution model and did not optimize any parameter, but all branch lengths were equal and set to the average of their original lengths. This strong model violation was used to measure the impact on ACR of branch lengths estimated from molecular data and possibly poorly fitted to the reconstruction of non-molecular characters.
- F81 model and rescaled branch lengths: we used the F81 model for DNA and F81-like model (see above) for proteins. All parameters (equilibrium frequencies, global rate p, smoothing parameter ε) were estimated from the true tree, which was rescaled accordingly before performing ACR. This setting corresponds to the standard, default option of PastML. The goal was to check that the loss of accuracy was low, compared to the perfect ‘true model and branch length’ setting.
- F81 model and flattened branch lengths: we combined the F81 (DNA) and F81-like (protein) models and optimizations, with flattened branch lengths, as described above. PastML was launched with the flattened tree and the F81 option described above. The goal was to measure the additional loss of accuracy when the branch lengths are ignored, compared to the previous setting.
- JC model and rescaled branch lengths: this setting is the same as ‘F81 model and rescaled branch lengths’, but using JC (DNA) and JC-like (proteins) models, which assume that all equilibrium frequencies are equal, as well as all substitution rates. The goal was to measure the loss of accuracy, compared to F81 and F81-like models.
- JC model and flattened branch lengths: we used JC and JC-like models with flattened branch lengths, thus a very strong model violation, which was used to measure the combined impact of ignoring both the substitution model and branch lengths. Parsimony is based on similar assumptions, but performs very different calculations.

### Comparison criteria

To compare the accuracy of the various ACR methods being tested, we used the Brier score of the predicted states against the known, true scenario. In the above Brier score formula *Truth* (*N, i*) was equal to 1 when the true state of node *N* was i, and 0 otherwise. Accordingly: *PPr* (*N, i*) was equal to 1 (*i* is predicted) or 0 (*i* is not predicted) for the methods predicting a unique state (i.e. Joint and MAP); *PPr*(*N,i*) was equal to 1*/k (i* is predicted) or 0 (i is not predicted) with Parsimony and MPPA when *k* states were predicted; with Marginal, *PPr* (*N, i*) was simply equal to the marginal posterior probability of state *i* for node *N*. The Brier scores of the nodes were then averaged, and we returned the average score over 2,500 trials (50 trees x 50 sites) for each of the simulation conditions.

To compare the performance of the various methods in producing consistent predictions across the tree, we applied the Brier score to edges instead of nodes. The goal was to check that the predictions for the two extremities of any given edge were compatible and close to the truth, thus establishing, or not, the superiority of global predictions as produced by Joint, over independent predictions as produced by MAP or MPPA (see also Gascuel and Steel 2014). An edge *E* was perfectly predicted (*PPr* (*E, i, j*) = *Truth* (*E, i, j*) = 1) whenitstwo extremity states *i, j* were the same as the true ones. In case of multiple predictions, *k* on one extremity and *p* on the other, *PPr* (*E, i, j*) was equal to 1/*kp* when the true states were included in the predicted states at both edge extremities, and 0 otherwise. With Marginal we simply used for *PPr* (*E, i, j*) the product of the state posteriors of *i* and *j* at both extremities of *E*. The formula to compute the Brier score was the same as for nodes, but considering a state space of size *s*^2^.

Lastly, for Parsimony and MPPA we counted the average number of predicted states per node in the various simulation conditions. The goal was to check the advantage of predicting several states instead of a single one, in case of uncertainty.

### Accuracy of the various ACR approaches

Brier score results are displayed in Figure 2 (four states, DNA-like) and Figure 3 (twenty states, protein-like) for the most relevant simulation conditions. In Table 1 we provide the number of predicted states for Parsimony and MPPA. Additional results are provided in Supplementary Material (Fig. S1, S2, S3) for all simulation conditions and the edge Brier score.

**Figure 2:**
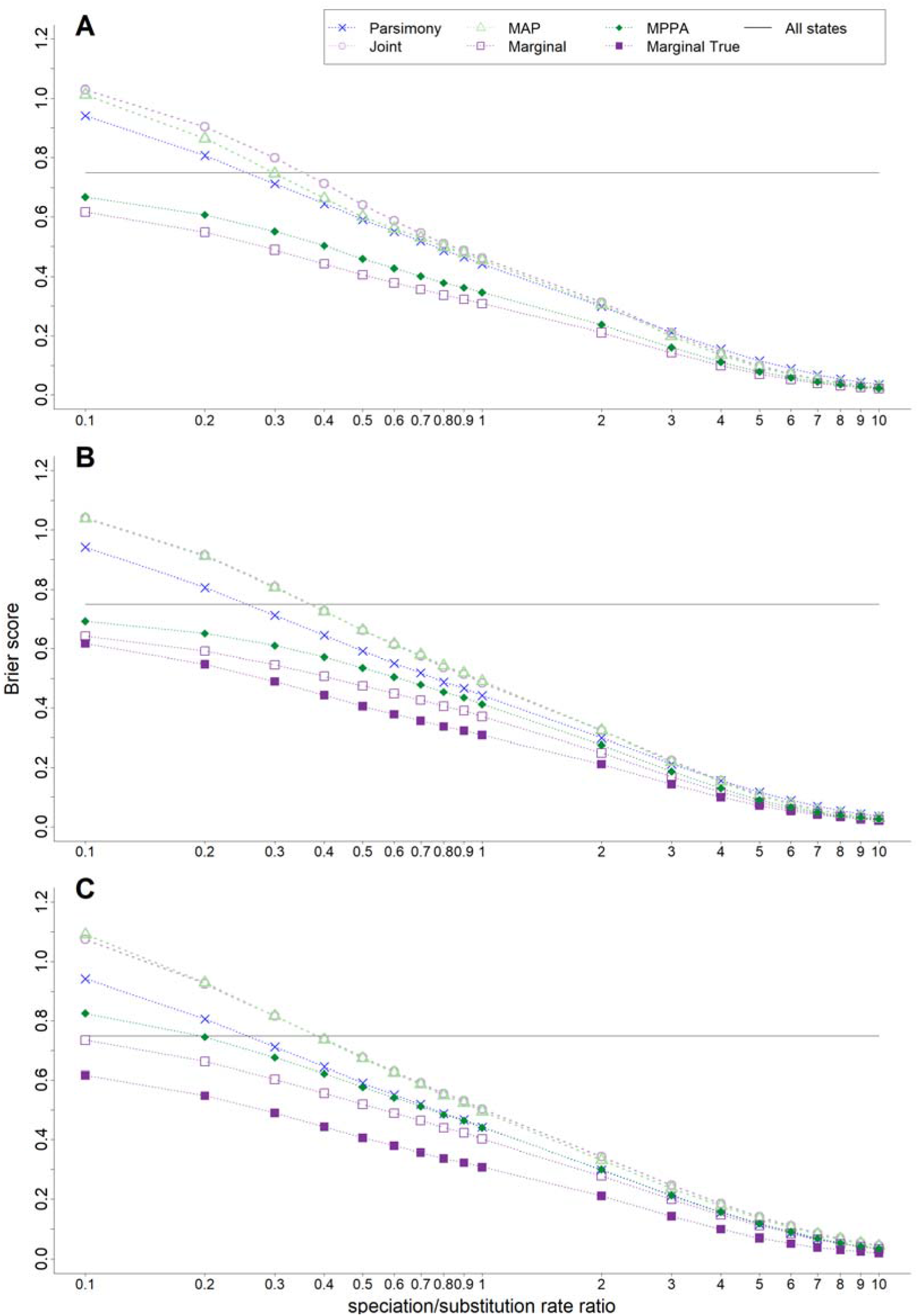
Accuracy of ACR methods with DNA-like (4 states) simulated data. X-axis: speciation/mutation rate ratio. Y-axis: Brier score. ‘All states’: all states are predicted with equal probability (= 1/4). ‘Marginal True’: best possible accuracy, obtained with Marginal using the mutation model and tree (including branch lengths) used to generate the data. ACR was performed with each ML method based on: A: the true model (HKY) and true branch lengths; B: the F81 model and rescaled branch lengths (default PastML option); and C: the JC model and flattened branch lengths (see text for details).

**Figure 3:**
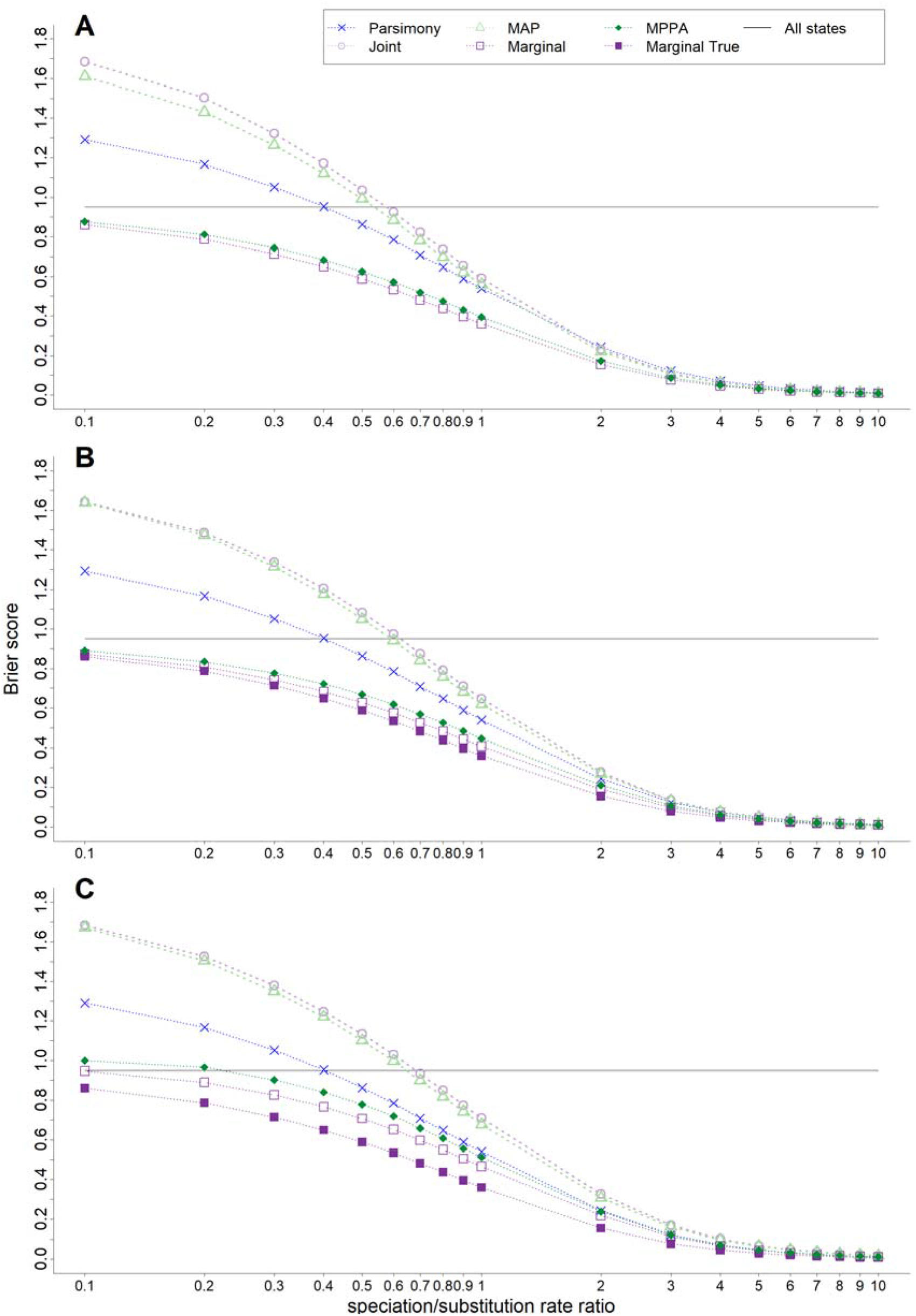
Accuracy of ACR methods with protein-like (20 states) simulated data. See note to Figure 2 and text for details. With ‘All states’ every state is now predicted with probability 1/20.

**Table 1.** Average number of states predicted per node by Parsimony and MPPA. HKY and true BL: the HKY model and the true branch lengths were used for ACR; F81 and rescaled BL: the F81 model was used and all parameters including the global rate and smoothing parameter were estimated from the data; then, the tree was rescaled accordingly and ACR was performed. JC and flat BL: the JC model was used and all parameters were estimated from the data using the tree with flattened branch lengths; then, the tree was rescaled accordingly and ACR was performed. JTT and true BL: same as ‘HKY and true’ but using JTT; F81-like and rescaled BL: same as ‘F81 and rescaled BL’, but using an F81-like model with 20 states (amino-acids). JC-like and flat BL: same as ‘JC and flat BL’ but using a JC-like model with 20 states. See text for details.

| speciation/<br>substitution | DNA-like data |  |  |  | Protein-like data |  |  |  |
| --- | --- | --- | --- | --- | --- | --- | --- | --- |
|  | Parsimony | HKY<br>true BL | F81<br>rescaled BL | JC<br>flat BL | Parsimony | JTT<br>true BL | F81-like<br>rescaled BL | JC-like<br>flat BL |
| 0.1 | 1.36 | 2.75 | 2.66 | 1.82 | 2.93 | 12.81 | 13.65 | 10.45 |
| 0.2 | 1.32 | 2.50 | 2.37 | 1.55 | 2.77 | 10.16 | 11.20 | 5.56 |
| 0.3 | 1.28 | 2.25 | 2.10 | 1.42 | 2.62 | 7.96 | 9.04 | 3.59 |
| 0.4 | 1.25 | 2.04 | 1.88 | 1.34 | 2.48 | 6.33 | 7.35 | 2.89 |
| 0.5 | 1.23 | 1.85 | 1.67 | 1.28 | 2.35 | 5.00 | 5.88 | 2.47 |
| 0.6 | 1.21 | 1.71 | 1.53 | 1.24 | 2.23 | 4.08 | 4.80 | 2.21 |
| 0.7 | 1.19 | 1.61 | 1.42 | 1.21 | 2.12 | 3.35 | 3.87 | 2.02 |
| 0.8 | 1.18 | 1.54 | 1.34 | 1.19 | 2.02 | 2.82 | 3.17 | 1.87 |
| 0.9 | 1.17 | 1.49 | 1.30 | 1.17 | 1.93 | 2.45 | 2.68 | 1.76 |
| 1 | 1.16 | 1.45 | 1.27 | 1.16 | 1.85 | 2.18 | 2.33 | 1.67 |
| 2 | 1.10 | 1.25 | 1.16 | 1.10 | 1.39 | 1.26 | 1.28 | 1.26 |
| 3 | 1.08 | 1.15 | 1.11 | 1.07 | 1.20 | 1.10 | 1.12 | 1.14 |
| 4 | 1.07 | 1.09 | 1.08 | 1.06 | 1.12 | 1.05 | 1.07 | 1.09 |
| 5 | 1.05 | 1.06 | 1.06 | 1.05 | 1.08 | 1.03 | 1.04 | 1.06 |
| 6 | 1.04 | 1.04 | 1.04 | 1.04 | 1.06 | 1.02 | 1.03 | 1.04 |
| 7 | 1.04 | 1.03 | 1.03 | 1.03 | 1.04 | 1.01 | 1.02 | 1.03 |
| 8 | 1.03 | 1.02 | 1.02 | 1.03 | 1.03 | 1.01 | 1.02 | 1.03 |
| 9 | 1.03 | 1.02 | 1.02 | 1.02 | 1.03 | 1.01 | 1.01 | 1.02 |
| 10 | 1.02 | 1.01 | 1.02 | 1.02 | 1.02 | 1.01 | 1.01 | 1.02 |

We observe that:

- As expected, predictions are very difficult with the lowest speciation/mutation rate ratios (0.1); then, all methods have similar or worse accuracy as/than the agnostic method predicting all states with equal probability. With higher speciation/mutation rate ratios, predictions become easy, as very few mutations occur in the tree. With the highest ratios (≥5) all methods succeed (Brier score ≈ 0) and are equivalent.
- As expected again, predictions are more difficult with 20 states (Fig. 3, Brier score ≈ 1.7 for the worst methods and conditions) than with 4 states (Fig. 2, Brier score ≈ 1.1 for the worse methods and conditions). Moreover, the gap between the best and worse methods is larger with 20 states than with 4 states. However, the ranking of the various methods is just the same in both settings.
- Marginal is the best method, as expected from decision theory, and its advantage still holds with strong model violations (Panels C in Figs. 2–3, JC-like model with flattened branch lengths). Joint and MAP are the worst, due to the fact that they predict a unique state and do not account for uncertainty. Their accuracies are similar, with a slight advantage for MAP in certain conditions (e.g. in the absence of model violations, Panels A in Fig. 2–3). This result still holds with the edge Brier score (Fig. S1, S2), thus indicating again (Gascuel and Steel 2014) that Joint and global predictions have no advantage compared to the more local calculations implemented in Marginal. Moreover (Fig. S1 to S3), all results obtained with the edge Brier score are congruent with those of the node Brier score, and the various model violations described above confirm the findings of Figures 2–3.
- Thanks to multiple predictions in uncertain configurations, Parsimony has a clear advantage over Joint and MAP (Tab. 1). The advantage of MPPA is even larger, due to the fact that MPPA predicts more states than Parsimony, and that these states are predicted using a rigorous probabilistic procedure. With medium speciation/mutation rate (1.0), the number of predicted state by MPPA is ~1.25 and ~2.5, for 4 and 20 states, respectively (Tab. 1). This indicates that the large accuracy gain of MPPA, compared to unique state prediction methods (Joint, MAP), is obtained thanks to a relatively low number of predicted states, which eases the interpretability and visualization of the global scenarios returned by MPPA. We shall see that these findings are confirmed with real HIV data.
- In terms of accuracy, MPPA is close to Marginal in all conditions and is the second best method. When violations occur in the mutation model (i.e. F81 and F81-like models are used to analyze data simulated with HKY and JTT, respectively, Fig. 2B and 3B; see Fig. S3 for JC and JC-like analyzes) MPPA accuracy remains close to the best possibly results that are obtained with Marginal using the true model. This advocates the use of F81-like model as default option, a choice which is not only computationally simple but also fairly accurate. With flattened trees, the accuracy gap between MPPA and Marginal using the true model becomes more substantial. Interestingly, with flattened branch lengths the number of states predicted by MPPA is close to the one predicted by Parsimony (both are based on similar assumptions), and significantly lower than the number of states predicted by MPPA with full branch lengths. With flattened branch lengths the uncertain and conflictual parts of the tree (e.g. nodes surrounded by long branches, tips with different states separated by short branches, etc.) are hidden, and less states are predicted. This indicates the importance of having (even approximate) branch lengths to obtain accurate state predictions.

To summarize, MPPA performs well in this simulation study, with accuracy close to the fully probabilistic Marginal method, but outputs that are much easier to interpret and visualize. Moreover, the F81-like model seems to be a relevant choice, as it yields accurate ancestral predictions while avoiding difficult estimations of the relative rates of change from one state to another.

## Results: Application to a Large HIV-1M Subtype C Data Set

### Data and Analyses

To demonstrate PastML’s performance on real data, we reconstructed the ancestral history of HIV-1M subtype C (HIV-1C) epidemics. We used a large data sets of 3,619 HIV-1C *pol* sequences, obtained from (Jung et al. 2012), (Chevenet et al. 2013), and the latest (2017) *pol* alignment of the Los Alamos HIV database (http://www.hiv.lanl.gov). This dataset is annotated with sampling dates and countries grouped into 11 regions (Central Africa, South Africa, Southern Africa excluding South Africa, West Africa, East Africa, Horn of Africa, North America, Central America, South America, Europe, and Asia; see Chevenet et al. 2013 for details). We built a PhyML tree (Guindon et al. 2010) from the DNA sequences, and rooted this tree using non-C sequences. To check the robustness of PastML inferences against phylogenetic uncertainty, the tree building and rooting procedure was repeated 5 times with different starting trees, resulting in 5 trees with clearly different topologies. We also checked the robustness of the results regarding state sampling variations, as some regions were sampled more intensively than others (e.g. 991 from Central Africa, and 64 from West Africa). For this purpose, we randomly pruned the tree by keeping at most 250 tips per region (i.e. 250 from Central Africa, and still 64 from West Africa). This severe pruning was repeated 5 times.

We reconstructed the phylogeography of HIV-1C epidemics using PastML. The historical worldwide diffusion of HIV-1C has been studied in several articles (Vidal et al. 2000, Hemelaar 2012, Faria et al. 2014, see also further references about subtype C sub-epidemics) and constitutes a good test bench for phylogeography methods.

PastML was also used to reconstruct the ancestral scenarios describing the emergence and diffusion, and reversion in some cases, of surveillance drug resistance mutations (SDRMs, Bennett et al. 2009). SDRMs emerge under the pressure of drug treatments, and then may be transmitted to drug naïve patients. Thus, an essential public health issue is to detect potential drug resistant sub-epidemics, which could become prevalent and pose major problems, as is already the case with other pathogens and diseases (e.g. malaria). Parsimony-based ancestral reconstructions were already used fruitfully in this context (Mourad et al. 2015, Zhukova et al. 2017), considering two character states: the SDRM is absent or present, and the corresponding strain (tip, node) is respectively sensitive or resistant. Five of the most prevalent SDRMs were analyzed by PastML with default options, and we performed analyses through time to study the dynamics of SDRM emergence, diffusion and reversion.

All these data are available from https://pastml.pasteur.fr/. See Material and Methods for details and method options. PastML analyses were performed on a laptop with a 4-core 2.50GHz CPU. ACR and visualization of HIV-1C phylogeography (11 states) took ~5 minutes per tree, ACR and visualization of SDRM dynamics (2 states) took ~10 seconds per tree.

### Phylo-geographic analyses

PastML results using MPPA are displayed in Figure 4, showing that the epidemic started in Central Africa. This agrees with the results of Faria et al. (2014), who showed using a Bayesian approach that HIV-1 subtype C originated in Mbuji-Mayi (Democratic Republic of Congo, DRC), in Central Africa, developed in the DRC mining regions and spread from there south and east, probably through migrant labor (see also Vidal et al. 2000 for the origin of HIV-1M). Our reconstruction shows several introductions of HIV-1C from Central Africa to Southern (violet) and South Africa (orange), forming infected clusters of different sizes. For instance there are 130 independent introductions to Southern Africa leading to small clusters of 1-8 patients, and three larger clusters of 36-79 patients and the largest South African cluster of 194 patients. Our reconstruction also shows 42 independent cases of HIV-1C spread from

**Figure 4:**
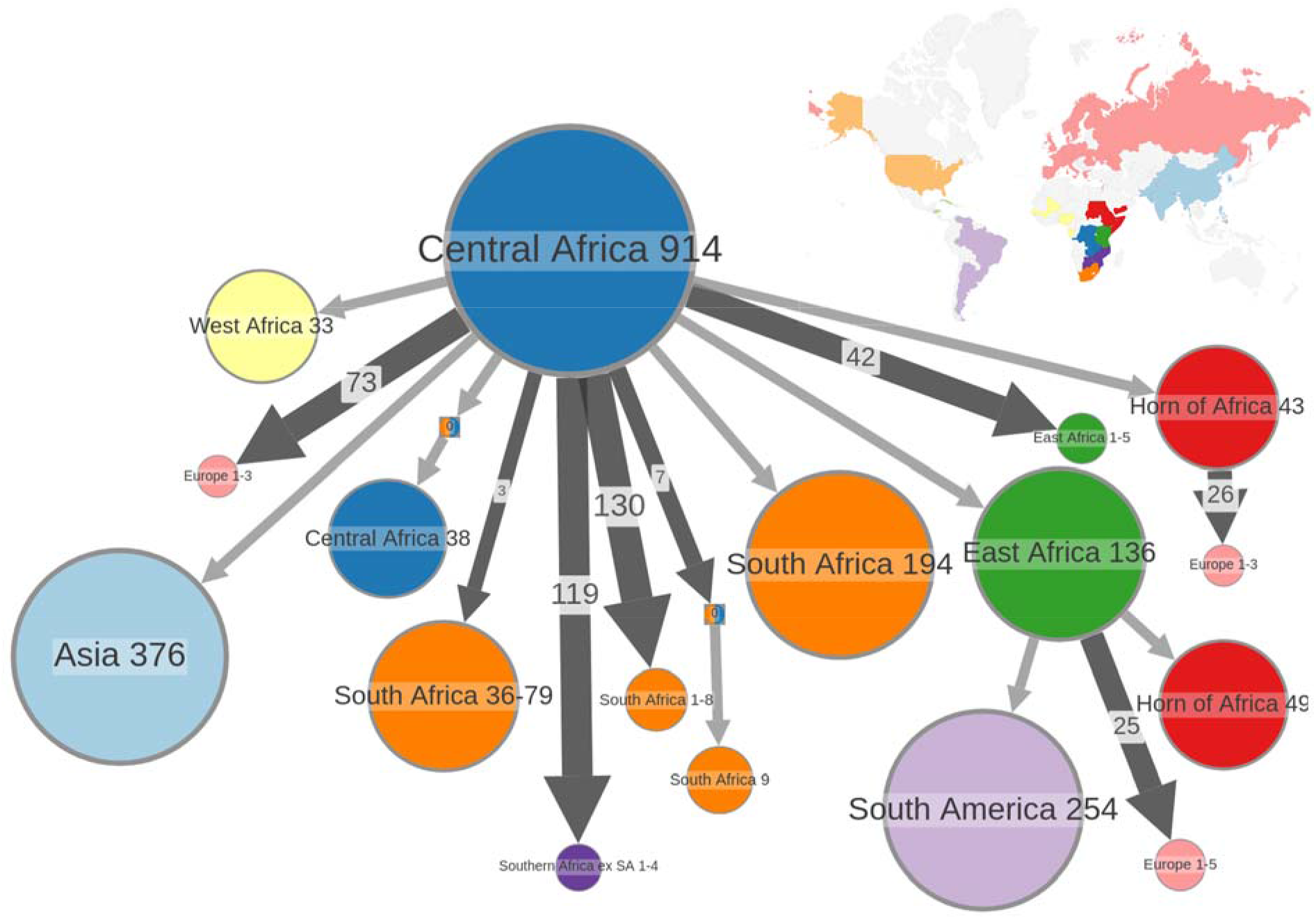
Ancestral reconstruction of HIV-1C epidemic locations. The figure shows the compressed visualization of the MPPA+F81 location reconstruction for HIV-1C, with minor details removed (10 as trimming threshold). Different colors correspond to different geographical regions as shown in the map in the top right corner.

Central Africa to East Africa that formed small clusters of 1-5 patients, and a larger one comprising 136 patients. From the latter one it was further introduced to South America (lilac cluster of size 254), and the Horn of Africa (red clusters of size 49). There is a second direct introduction to the Horn of Africa from Central Africa (red cluster of size 43 directly connected to the root cluster), which could be a plausible explanation for the observation of the C and C’ sub-clusters in Ethiopia (Abebe et al. 2000; see also Chevenet et al. 2013). Lastly, we recover the result of Jung et al. (2012), showing a Central Africa origin of the HIV-1C epidemic in West Africa, especially in the Senegalese MSM population.

The study of Soares et al. (2003) has shown evidence that subtype C has entered Brazil as a single introduction, or at least as a very small group of genetically related viruses. Our reconstruction confirms this statement: in our dataset 245 out of 254 South American sequences come from Brazil, and we can see 244 of them in the lilac South American cluster of size 254.

Our reconstruction also shows a major introduction of HIV-1C epidemic from Central Africa to Asia forming a large cluster of 376 patients (light-blue). Most of the Asian sequences in our dataset (371 out of 404) come from India, and it was previously shown that the Indian HIV-1C epidemic originated from a single or few genetically related African lineages (Neogi et al. 2012). The introductions to Europe, on the other hand, are multiple, from different regions and lead to smaller clusters (in pink), a finding already pointed out in numerous studies.

Figure S4 in Supplementary Material indicates that these finding are remarkably robust against phylogenetic uncertainty. While the 5 trees being tested are quite different (topological distance >30%, see Material and Methods), the main components (as shown in Fig. 4) of the ancestral scenarios reconstructed using these trees are nearly identical. The same holds with the sampling rate (Fig. S5), where our severe pruning of the most represented locations (>250 tips, see above) does not change substantially the results.

### Dynamics of SDRM emergence, diffusion, and reversion

Results for M184V, which is the most prevalent SDRM in our data set, and the combination with phylogeography for the largest resistant cluster are displayed in Figure 5. M184V is a major non-nucleoside RT inhibitor (NRTI) mutation selected in patients receiving Lamivudine (3TC) and Emtricitabine (FTC) (Gallant 2006). 3TC was approved for medical use in the United States in 1995, and FTC in 2006. They are both used worldwide nowadays. According to the study of Castro et al. (2013) on the persistence of SDRMs in the absence of drug-selective pressure, M184V median time to loss is 1 (0.5-2.0) years.

**Figure 5:**
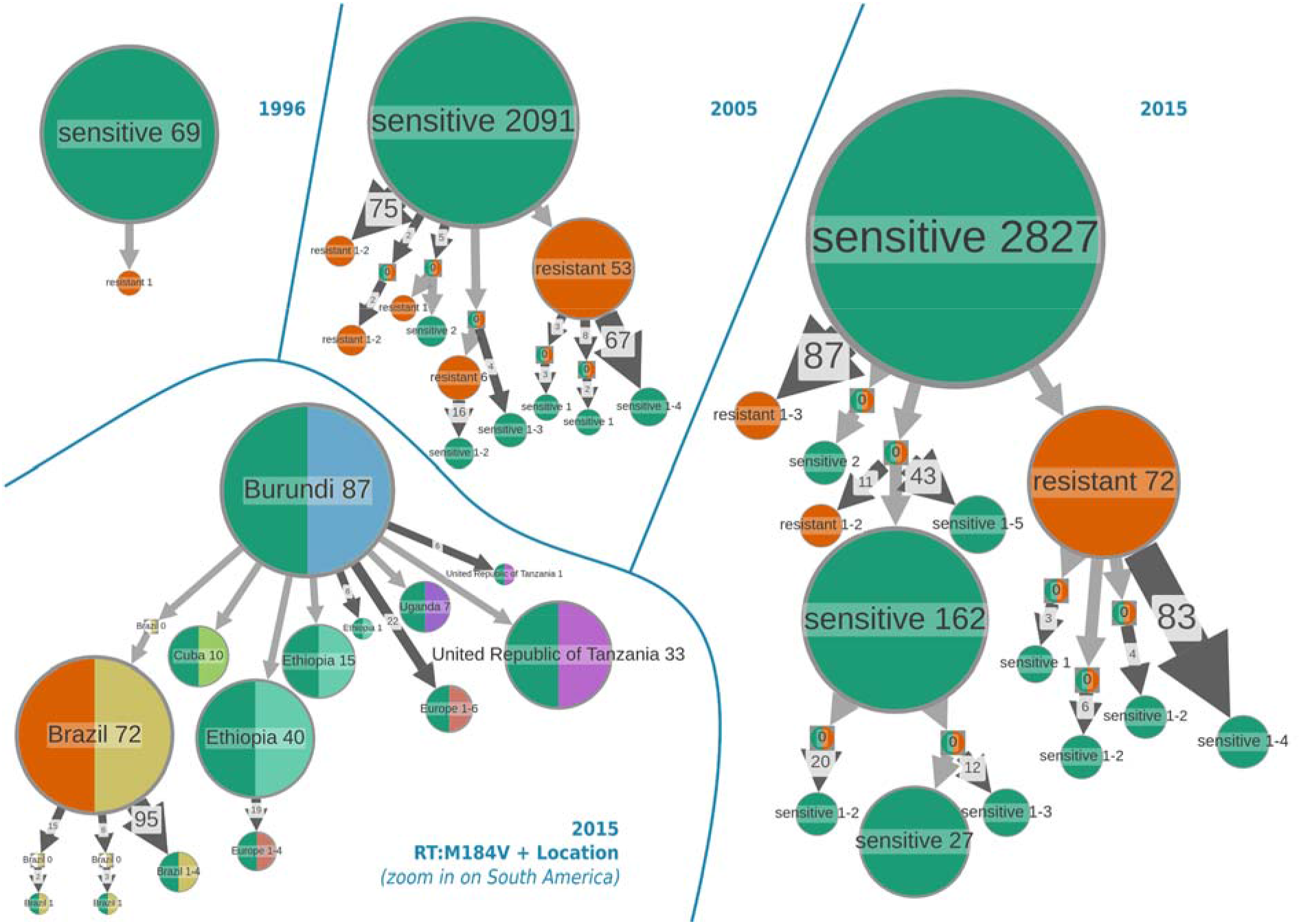
Ancestral state reconstruction of the presence/absence of DRM M184V over time (top), combined with location data (bottom left). The reconstruction was done with MPPA+F81. For the timeline the tree was pruned at each year to remove the tips sampled after that year prior to the reconstruction. In the bottom left panel M184V presence/absence is combined with location data: M184V state is shown color-coded in the left half of each node (green when mutation is absent, and orange for resistant strains), countries are color-coded in the right half of each node, and shown in the labels.

Ancestral state reconstruction allows us to detect potentially acquired (ADR) and transmitted drug resistance (TDR) patterns. An acquired drug resistance is represented by a single-patient resistant node in the compressed visualization, which implies a state change from a sensitive parent node. Potential TDRs are represented by cluster(s) of resistant patients, where internal edges correspond to transmissions of resistant strains. Note that these simple statements still hold with incomplete sampling (Mourad et al. 2015): transmissions of SRDMs within resistant clusters are then indirect, while a one-patient resistant node may correspond to a small resistance cluster, the root of which acquired the DRM.

We analyzed the reconstructed transmission tree at different time points, each time pruning the tree to remove the nodes sampled after the corresponding year. Figure 5 shows the results for 1996 (the sampling year corresponding to the first sequences with M184V in our dataset), 2005 and 2015 (last sampling year in our dataset). We see the appearance and growth of potential TDR clusters over time. In 2005 the main configurations included a major TDR cluster of 53 patients, and 78 cases of independent DRM emergence (ADR, i.e. one-patient node, or small resistant clusters of two patients). There are also multiple cases of reversion of the DRM (e.g, 67 cases of patients having a sensitive virus at the time of sampling, which originate from the major resistant cluster). By 2015 the main TDR cluster grew (to 72 patients), and so did the numbers of cases of ADR and DRM reversion. Importantly, the main resistance cluster in 2005 is included in the one of 2015, which demonstrates the potential of the method in surveying the emergence of problematic resistant sub-epidemics, as the 2015 cluster was already predictable in 2005.

To further investigate the largest TDR cluster we combined the ancestral state reconstruction for M184V with the location. The result located the whole resistant cluster in South America. We then increased the geographical resolution by replacing the regions with the countries and focused on the subtree with the root in East Africa, as this is from where the virus was spread to South America according to our reconstruction (Fig. 4). The results are shown in the bottom left panel of Figure 5, and suggest that the resistant cluster is located in Brazil, and that it originated from either a sensitive or a resistant case in Brazil (the parent node of the TDR cluster is a Brazilian node with unresolved M184V state). It also shows that the virus was introduced to Brazil from Burundi, from where it was also spread to Tanzania, and Ethiopia. This geographical result agrees with the study on HIV-1C epidemics in Eastern Africa and Southern Brazil by Mir et al. (2018), although the latter was performed using a Bayesian approach and a different dataset, which included more Brazilian sequences.

The results for the 2nd, 3rd and 4th most prevalent SDRMs in our dataset (K103N, D67N, and K70R) are similar to those for M184V: they show emergence and growth of TDR clusters over time, and well as growth of the number of ADR and reversions to sensitive state. The largest TDR cluster is located in Brazil. With the decrease of SDRM prevalence, the size of TDR clusters in our data decreases, from a 72-patient TDR cluster for M184V to a 7-patient one for K70R. The analysis of K103N can be found in Supplementary Material (Figure S6). The results for the 5th most prevalent SDRM (Y181C) are different: We hardly see any TDR clusters (their size is at most 4 patients), and the largest TDR cluster is located in India (see Figure S7). This could be the very start of TDR spread for this mutation hence making it a candidate for closer surveillance, or it could simply be due to Y181C’s quick reversion time (median of 1.3 years, cf. Castro et al. 2013) and hence inability to form TDR clusters.

## Conclusions

We presented a new, simple approach to reconstruct ancestral scenarios, deal with the uncertainty of ancestral inferences in the difficult regions of the tree (typically around the tree root), and visualize and edit interactively the tree-shaped graphical representation of the most likely ancestral scenarios. All the results obtained with a large HIV-1 subtype C dataset are fully congruent with previous studies, and are obtained very quickly using a simple laptop. Moreover, these results are robust against phylogenetic uncertainty and sampling rate variations. This study on HIV-1C (and others in preparation) indicate that the signal extracted by ACR is remarkably strong and bears a very useful information in a molecular epidemiology context.

Directions for further research include the development of methods and tools to compare our tree-shaped, compressed representations of ancestral scenarios, calculate some distance between two scenarios, extract the common parts and the differences, and propose some consensus. Moreover, the current version of PastML is based on JC-like and F81-like models. Some refinement (e.g. in the line of Lemey et al. 2014, and Dudas et al. 2017) should be useful, not only to improve the accuracy and ancestral reconstructions, but also to provide users with a global view of the evolutionary processes at stake (strain flow between regions and countries, acquisitions and losses of molecular characters, dynamics of ecological character changes, etc.).

## Material and Methods

### Computation of the Marginal Posterior Probabilities

Let *N* be a given internal node of *T*, and *U*, *V* and *F* be the left descendant, right descendant and father of *N*, respectively, with corresponding rescaled branch lengths denoted as *u, v* and *f.* Moreover, let *Down* (*N*) be the vector of state conditional likelihoods induced by the state values of the tips of the “*down*” subtree rooted with *N*. *Down* (*N, i*) is the ith component of *Down* (*N*). *Down* (*N, i*) is equal to the likelihood of having state *i* in *N* given the states observed in the extant descendants of *N*.

*Down* (*N*) is computed recursively using the pruning algorithm (Felsenstein 1981), which combines a post-order tree traversal with the following formula:

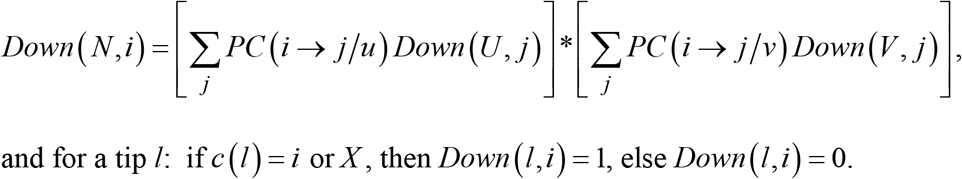

This algorithm proceeds in a bottom-up fashion, first computing the conditional likelihoods of the nodes close to the tips and progressing until the tree root (e.g. we first compute *Down* (*U*) and *Down* (*V*), and then *Down* (*N*)). The conditional likelihoods so obtained can be used to compute marginal posterior probabilities and then predict the ancestral states attached to every tree node. Several ancestral reconstruction programs use this approach. However, a more accurate method does exist (Yang 2007). Indeed, when using *Down* (*N*) we only account for the information contained in the tips descending from *N*, and not for the information contained in the rest of the tree.

To account for all tree information, we define a second vector of conditional likelihoods attached to *N*, *Up* (*N*), where *Up* (*N, i*) denotes the conditional likelihood of having *i* in *N* given the tip values observed in the “*up*” subtree of *N*. To define this subtree, assume that *T* is re-rooted with N; then *N* has three direct descendants: *U, V* and *F*, each associated to a subtree. The *up* subtree of *N* is defined as the subtree associated to *F* including the branch (of length *f* from *F* to *N.* In other words the *up* subtree contains all branches, nodes and tips which are not included in the *down* subtree of *N.* To compute the *Up* conditional likelihoods we use formula (applied to node *U* to simplify the notation, but the same formula applies to *V, N* and all tree nodes):

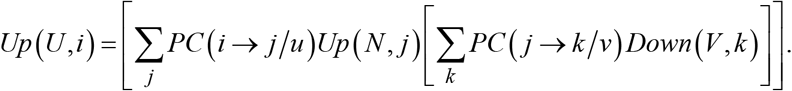

This formula is exploited recursively thanks to a top-down, pre-order tree traversal. We start from the tree root *R*, having *Up* (*R, i*) = 1 for all states i, and progress toward the tips; for example, *Up* (*N*) is computed after *Up* (*F*) and before *Up* (*U*) and *Up* (*V*), as seen in the formula. Moreover, this formula uses the *Down* likelihoods, which have to be computed first. Both *Down* and *Up* calculations are easily extended to polytomies: the *Down* formula contains as many sum terms as *N* has descendants (instead of 2 above with *U* and V); the *Up* formula contains as many internal sum terms as *U* has brother nodes (instead of 1 above with *V*).

Once *Down* (*N*) and *Up* (*N*) have been computed, the state marginal posterior probabilities of *N* are computed using (remember that for the tree root *Up* (*R, i*) = 1):

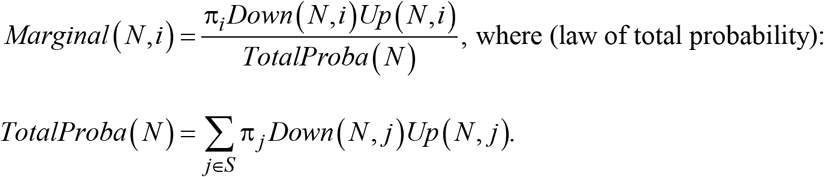

These algorithms have a time complexity in *O* (*ns*^2^), where *n* is the number of tree tips and s the number of states. The whole procedure is thus linear in *n* and remarkably fast.

All along these calculations, some of the conditional likelihood values may be extremely small when *n* is large, and smaller than the minimum value permitted for double floating numbers. As in other ML programs, if the conditional likelihoods of *N* are smaller than a given threshold, then all conditional likelihoods of *N* are multiplied by a power of 2. This numerical trick does not change the marginal posterior probabilities, as the relative values of the conditional likelihoods are preserved.

### HIV-1C data and analyses

We used an HIV-1C *pol* sequence dataset, previously used in (Jung et al. 2012), and then (in a slightly updated version) in (Chevenet et al. 2013). We extended the latter alignment with HIV-1C *pol* sequences from the latest (2017) *pol* alignment in the Los Alamos HIV database (https://www.hiv.lanl.gov/content/index), hence adding 583 sequences not present in Chevenet et al. (2013). Addition of the new sequences was performed using MAFFT multiple sequence alignment program with the --add option (Katoh et al. 2013). The final alignment contains 3,619 HIV-1C *pol* sequences, plus 35 outgroup reference sequences from the non-C subtypes. The dataset is annotated with sampling dates and countries grouped into 11 regions: 12 sequences from North America, 26 from Central America, 256 from South America, 366 from Europe, 404 from Asia, 64 from West Africa, 144 from the Horn of Africa, 991 from Central Africa, 224 from East Africa, 353 from Southern Africa excluding South Africa, and 777 from South Africa (see Chevenet et al. 2013 for details).

We detected the Surveillance Drug Resistance Mutations (SDRMs) in the alignment, using the Sierra web service of the Stanford HIV drug resistance database (Liu et al. 2006). We removed the SDRM positions from the alignment and reconstructed 5 most parsimonious trees using TNT (Goloboff et al. 2016), which were used as starting trees for 5 runs of PhyML (Guindon et al. 2010) with GTR+I+Γ6 substitution model and aLRT SH-like branch supports. The resulting trees were rooted with the outgroup sequences, which were subsequently removed from the trees. The branches of length zero and aLRT SH-like support less than 0.5 were collapsed into polytomies. We thereby obtained 5 ML trees with clearly different topologies. The average normalized bipartition distance was equal to 0.33 (ETE 3 toolkit, Huerta-Cepas et al. 2016), and the average quartet distance to 0.31 (tqDist library, Sand et al. 2014), where 0.0 means identical trees, and 1.0 corresponds to trees that have no bipartition / no quartet in common. These multiple trees were used to check the robustness of our ancestral reconstructions against phylogenetic uncertainty. Results in Figures 3 and 4 are provided for the most likely tree. Results for all trees are in Supplementary Materials (Fig. S4, scenario 4 corresponds to the most likely tree used in Fig. 3, 4, and S5).

PastML was used with default options (MPPA prediction method, F81 model, and a trimming threshold of 10 to remove minor details) to reconstruct the ancestral locations of all tree nodes, among the 11 regions (character states) present in the dataset. We also checked the robustness of the results regarding state sampling variations, as some regions were sampled more intensively than others. For this purpose, we pruned the tree by keeping at most 250 tips per region (for the regions with less samples all the tips were kept, for those with more samples 250 random tips were kept) and performed ancestral state reconstruction of the location for 5 such trees. Results in Figure 4 are for complete sampling. Results for partial sampling are in Supplementary Material (Figure S5).

PastML was also used to reconstruct the ancestral scenarios describing the emergence and diffusion, and reversion in some cases, of surveillance drug resistance mutations (SDRMs, Bennett et al. 2009). We analyzed SDRMs with high prevalence in our dataset: M184V with a prevalence of 0.07 (highest prevalence in our dataset), K103N (prevalence = 0.05, second highest prevalence), and Y181C (prevalence = 0.03, fifth highest prevalence). Results (available on request) for the third and fourth highest prevalence SDRMs are similar to those of M184V and K103N. PastML was used with default options, and we performed analyses through time to study the dynamics of SDRM emergence, diffusion and reversion. In this context, we have two character states: the SDRM is absent or present, and the corresponding strain (tip, node) is sensitive or resistant, respectively. Results for the most prevalent SDRM (M184V) are provided in Figure 4. Results for the two other SDRMs are in Supplementary Materials (Fig. S6, S7).

All the data and pipelines used to reconstruct the trees and analyze them are available from https://pastml.pasteur.fr.

## Acknowledgements

Sincere thanks to L. Blassel, S. Cosentino, S. Duchêne, F. Lemoine, M. Matsui, H. Ménager, S. Mestack, and K. Theys for their help and comments. This work was supported by the EU-H2020 Virogenesis project (grant number 634650, OG), by the INCEPTION project (PIA/ANR-16-CONV-0005, AZ and OG), and by Postdoctoral Fellowship and KAKENHI 282725 (SI), 16H06279 (WI), and 16H06154 (WI) from Japan Society for the Promotion of Science.

